# Human-derived microRNA 21 regulates indole and L-tryptophan biosynthesis transcripts in a prominent gut symbiont

**DOI:** 10.1101/2024.08.15.608161

**Authors:** Kayla Flanagan, Kirsten Gassner, Michaela Lang, Jurgita Ozelyte, Bela Hausmann, Daniel Crepaz, Petra Pjevac, Christoph Gasche, David Berry, Cornelia Vesely, Fatima C. Pereira

## Abstract

In the gut, microRNAs (miRNAs) produced by intestinal epithelial cells are secreted into the lumen and can shape the composition and function of the gut microbiome. Crosstalk between gut microbes and the host plays a key role in irritable bowel syndrome (IBS) and inflammatory bowel diseases, yet little is known about how the miRNA-gut microbiome axis contributes to the pathogenesis of these conditions. In this study, we aimed to explore the ability of miR-21, a miRNA that we found decreased in stool samples from IBS patients, to associate with and regulate gut microbiome function. Incubation of human faecal microbiota with miR-21 revealed a rapid association with microbial cells, reproducible across multiple donor samples. Fluorescence-activated cell sorting and sequencing of microbial cells incubated with fluorescently-labelled miR-21 identified organisms belonging to the genera *Bacteroides*, *Limosilactobacillus*, *Ruminococcus*, or *Coprococcus* which predominantly interacted with miR-21. Surprisingly, these and other genera also interacted with a miRNA scramble control, suggesting that physical interaction and/or uptake of these miRNAs by gut microbiota is not sequence-dependent. Nevertheless, transcriptomic analysis of the gut commensal *Bacteroides thetaiotaomicron* revealed a miRNA sequence-specific effect on bacterial transcript levels. Supplementation of miR-21, but not of small RNA controls resulted in significantly altered levels of many cellular transcripts and increased transcription of a biosynthetic operon for indole and L-tryptophan, metabolites known to regulate host inflammation and colonic motility. Our study identifies a novel putative miR-21-dependent pathway of regulation of intestinal function through the gut microbiome with implications for gastrointestinal conditions.

## Introduction

Irritable bowel syndrome (IBS) is a common chronic functional gastrointestinal disorder with a global prevalence of 11%, defined by disturbances in bowel movement habits and abdominal pain that lacks obvious signs of inflammation (1). The gut microbiota has been linked to the development of IBS (2,3). A reduced gut microbiome diversity and the abundance of *Clostridiales*, *Prevotella*, and methanogenic species have been postulated as an IBS-specific microbiome signature linked to symptom severity (2). Transplantation of faecal matter from healthy donors can lead to a transient improvement of IBS symptoms (4). Like in IBS, microbiota is also implicated in the pathogenesis of inflammatory bowel disease (IBD) (5–7), a chronic inflammatory disease that affects 0.3% of the population in the industrialized world (8). IBD is thought to be the result of an interplay between the host’s genetic predisposition, the immune system, and various environmental factors (9–11). A typical feature of IBD is an unstable and less diverse intestinal microbial community (5–7), displaying an increased abundance of species with pro-inflammatory properties such as members of Enterobacteriaceae, and a decreased abundance of anaerobic commensals such as Bacteroidota and Bacillota (12–14). Many studies have shown that disruption of microbial metabolism, such as short-chain fatty acid production, is implicated in the pathogenesis of IBD (15, 16). Indeed, efforts to target microbial pathways, either by altering the gut microbiota or through novel small molecule drugs, are currently seen as promising approaches to improve IBD symptoms and gut inflammation (17). Despite recent improvements, biomarkers and potential therapeutic targets for both IBD and IBS remain scarce.

MicroRNAs (miRNAs) are short non-coding RNAs, 18-23 nucleotides in length, that regulate gene expression in eukaryotes by binding to the 3’-untranslated regions of target messenger RNAs (mRNAs) through partial sequence homology, resulting in decreased stability and translation repression (18). A single miRNA can target hundreds of mRNAs and modulate the expression of many genes. As a direct consequence, miRNAs can regulate key biological processes such as apoptosis, cell proliferation, and immune cell differentiation, and play an important role in the development of disease (19). MiRNAs have attracted increasing attention in gastrointestinal disorders because they target molecules in pathways that regulate the intestinal epithelial barrier, inflammation, and cell migration(20–22). The miRNA miR- 21-5p (miR-21) is perhaps the most studied miRNA in this context, and its dysregulation has been implicated in a variety of gastrointestinal disorders including IBD (23), IBS (24), and colorectal cancer (25). MiR-21 contributes to IBD pathophysiology in part through its ability to increase gut barrier permeability via targeting proteins involved in cytoskeleton organization and tight junction regulation (26, 27), as well as via inducing Th2 immune cell differentiation (28). Increased miR-21 expression in the inflamed gut tissue is positively associated with IBD activity status (29). In addition to tissue, some studies have reported an increase in stool miR-21 levels in IBD patients when compared to healthy controls (30, 31), although the consequences of this for IBD pathogenesis, if any, are unknown.

MiRNA secretion into the extracellular space is thought to be a regulated process that acts as a mode of communication between cells, tissues, and even kingdoms (32–34). In the gut, miRNAs produced by intestinal epithelial cells are normally secreted into the gut lumen where they encounter and potentially interact with the gut microbiota (35). Mice lacking intestinal miRNAs exhibit an altered gut microbiome and increased susceptibility to colonic inflammation (colitis), and transplantation of faecal miRNAs from wild-type animals restores the microbiome and ameliorates colitis (35, 36). However, the mechanistic details of this miRNA-microbiota interaction and how it may impact colitis and microbiota homeostasis during miRNA dysregulation have not been fully elucidated. In addition, a negative correlation was found between the presence and abundance of microbes and levels of intestinal miRNAs, suggesting a potential role for the gut microbiota in the degradation or uptake of luminal miRNAs (36, 37). Indeed, miRNAs such as hsa-miR-515-5p and hsa-miR-1226-6p co-localize with pure-culture cells of *Fusobacterium nucleatum* or *Escherichia coli,* respectively, and have been demonstrated to modulate bacterial transcript levels and growth in a miR-specific manner (35). Whether each miRNA targets single or multiple taxa within the complex gut microbiome, and which taxa preferentially uptake or interact with miRNAs remain to be investigated.

To further elucidate the roles of miRNAs in host-microbiota interactions, we sought to determine whether the faecal miRNA miR-21 interacts with the microbiota and regulates its function. In this study, we establish the dynamics of association between miR-21 and the human gut microbiota. Using fluorescently-labelled miRNAs, fluorescence-activated cell sorting, and downstream 16S rRNA gene amplicon sequencing, we identify bacterial taxa within faecal communities that physically interact with miR-21. We further demonstrate that miR-21 modulates transcript levels in pure cultures of the gut symbiont *Bacteroides thetaiotaomicron*, including genes involved in the production of L-tryptophan, whose altered intestinal metabolism has been linked to gastrointestinal dysfunction in IBD (37) and IBS (38). Collectively, our findings reveal a mechanistic basis for the miR-21-microbiome axis and its role in gastrointestinal disorders.

## Results

### miR-21 is decreased in stool samples from IBS patients and faecal microbiota promotes miR-21 depletion *ex vivo*

MiR-21 is important for normal gastrointestinal function and altered levels of miR-21 are found not only in tissue but also in the stool of IBD patients (31). However, it has not been investigated whether miR-21 is also dysregulated in IBS and contributes to IBS pathogenesis. Thus, we started by quantifying the relative levels of miR-21 in stool samples from a cohort of IBS patients (*n*=6) and controls (*n*=5) (**Table S1**). All IBS patients included had IBS of the mixed-type (IBS-M) (39). The control group included individuals with no symptoms of intestinal disease undergoing colorectal cancer screening with no pathological findings at colonoscopy. Detection and quantification of miR-21 by qPCR revealed significantly decreased levels of miR-21 in stool samples from IBS patients compared to the control group (**Fig. 1A, Table S2**). Levels of miR- 26b, a miRNA used in many studies as an endogenous or normalisation control due to its high stability (40), were not significantly altered (**Fig. 1A**). Thus, in our small cohort, IBS-M patients present lower levels of miR-21 in stool.

**Figure 1.**
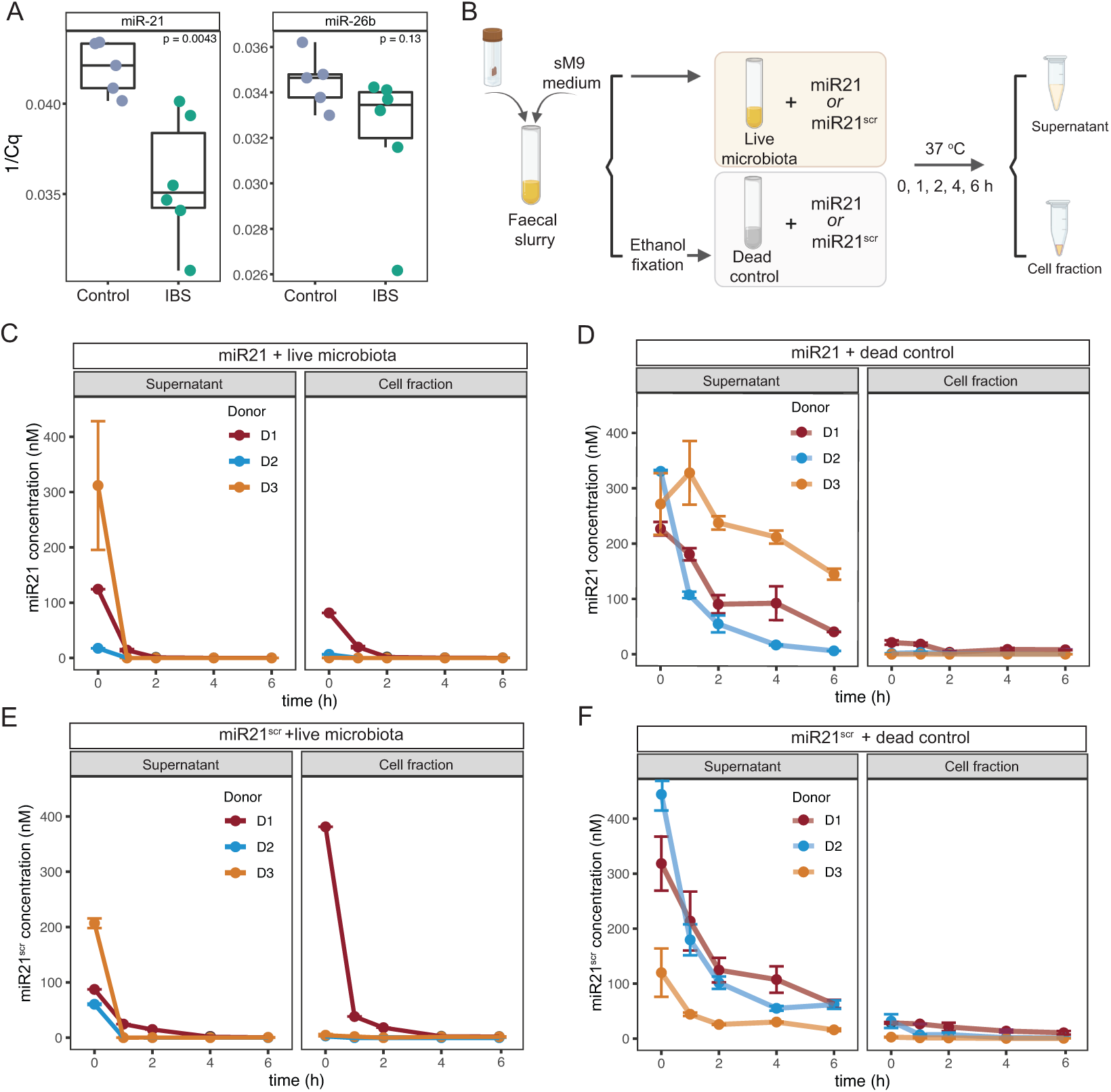
Quantification of miRNAs in faecal samples and dynamics of association of miRNAs with faecal microbiota. (A) Quantification of miR-21 and mir-26b in stool samples from colonoscopy controls (Control, *n*=5) and IBS-M patients (IBS, *n*=6) by qPCR. All samples were normalized to input amount of RNA. P-values were determined using an unpaired two-sample Wilcoxon test. **(B)** Schematic representation of miR-21 and miR-21 scramble control (miR-21^scr^) incubations with live or ethanol-fixed (dead control) microbiota derived from faecal samples. Each miR was added to a final concentration of 250 nM. Incubations were set in duplicates and sampled immediately after amendment (0 hours of incubation) or after 1, 2, 4 and 6 hours of incubation. Collected samples were centrifuged and further split into a supernatant (cell-free) and a cell pellet fraction (see Materials and methods) and processed for miRNA quantification by qPCR. **(C, D)** miRNA concentration over time in live (C) or dead control (D) microbiota incubations amended with miR-21, determined by qPCR. Three independent sets of incubations were established using faecal samples from three healthy donors (D1, D2, D3). **(E, F)** miRNA concentration over time in live (E) or dead control (F) incubations amended with miR-21^scr^, for the same three donors. Data points represent the mean of two replicates per condition and bars represent the standard deviation.

Several human-derived miRNAs have been shown to co-localize with nucleic acid signal (DAPI staining) in gut bacteria, which suggests an ability of miRNAs to enter into the cytoplasm of microbes (35, 41). To determine whether miR-21 can interact with microbes, we prepared faecal suspensions from samples collected from three different healthy donors and supplemented them with a synthetic miR-21 (**Fig. 1B**). We then followed the transfer of miR-21 from the supernatants into the microbial cells by sampling the vials at different time points post-supplementation (**Fig. 1B**). After separation into cell-rich (cell fraction) and cell-free (supernatant) fractions, levels of miR-21 in each fraction were quantified by qPCR (**Fig. 1B**). Results revealed a complete depletion of miR-21 in supernatant fractions from all donors within two hours of incubation (**Fig. 1C**, left panel). The addition of a ribonuclease (RNAse) inhibitor to the vials did not alter this pattern (**Fig. S1**), indicating that miRNA depletion is not due to the activity of common eukaryotic RNAses such as RNAse A. Due to the lack of a suitable bacterial RNAse inhibitor, it remains to be determined whether bacterial RNAses contribute to miR depletion. While miR-21 was detected in microbial cell fractions of all three donors at early time points (**Fig. 1C**, right panel, T0 and T1), it rapidly disappeared, indicating a fast but transient association with the microbiota.

To determine the potential contribution of active microbiota-independent processes to the depletion and/or degradation of miR-21 during incubations, dead controls using ethanol-fixed microbiota were established in parallel (**Fig. 1B,D**). Incubation of miR-21 with dead microbial cells showed a much slower decrease in miR-21 levels in supernatants over time for all three donors (**Fig. 1D**, left panel), with 25 ± 21% of the initial miR-21 remaining in supernatants at the end of the incubation. These values are considerably higher than the residual levels detected at the end timepoint for incubations with live microbiota (26.9E-04 ± 9.5E-04% of initial miR remaining after 6 hours) indicating that the faecal microbiota actively depletes miR-21. Importantly, miR- 21 levels remained stable in vials containing incubation medium in the absence of microbiota (**Fig. S2**). Small amounts of miR-21 were also detected in dead cell fractions, suggesting a degree of non-specific binding or adhesion of miR-21 to dead cells (**Fig. 1D**, right panel), which may contribute to its disappearance from supernatant fractions. Overall, our results suggest a rapid association of miR-21 with live microbial cells and a significant contribution from the live microbiota in depleting miR-21. The fact that we cannot detect the accumulation of miRNAs within cell fractions may reflect a short half-life of these molecules once inside microbial cells.

To investigate whether the association between miR-21 and gut microbiota is sequence-specific, additional incubations were set up using a miR-21 scramble control (miR-21^scr^), which contains the same nucleotides as miR-21 but arranged in a different order (**Fig.1B** and Materials and Methods). Quantification of miR-21^scr^ by qPCR in both supernatant and cell fractions revealed a rapid depletion of miR-21^scr^ from supernatants within two hours of supplementation (**Fig. 1E**, left panel). Similarly, to miR-21 incubations, miR-21^scr^ levels decreased more gradually in dead microbiota controls compared to live microbiota (**Fig. 1F**), with 16 ± 2.9% of the initial miR-21^scr^ remaining in supernatants at the end of the dead-control incubation, a significantly higher amount compared to live incubations (1.28E-03 ± 8.9E-04%). These results strongly suggest that the physical association and/or uptake of these miRNAs by gut microbiota is not sequence-dependent. The initial (T0 and T1) patterns of association of microbiota with miR-21 and miR-21^scr^ differed among donors, with higher levels of miR21 detected in cell fractions for donor 1 (**Fig. 1C, E**, right panels). These differences may be attributed to different microbiota compositions of the three donor samples (**Fig. 2B**). Nevertheless, the overall pattern of miRNA levels during incubations was similar across all three donors, suggesting that the association dynamics of these miRs with microbiota in this cohort is donor-independent. Thus, we conclude that miRNAs associate with and/or are internalized by faecal microbial cells. This association and subsequent depletion are independent of the miR sequence and appear to be independent of faecal microbiota composition.

**Figure 2.**
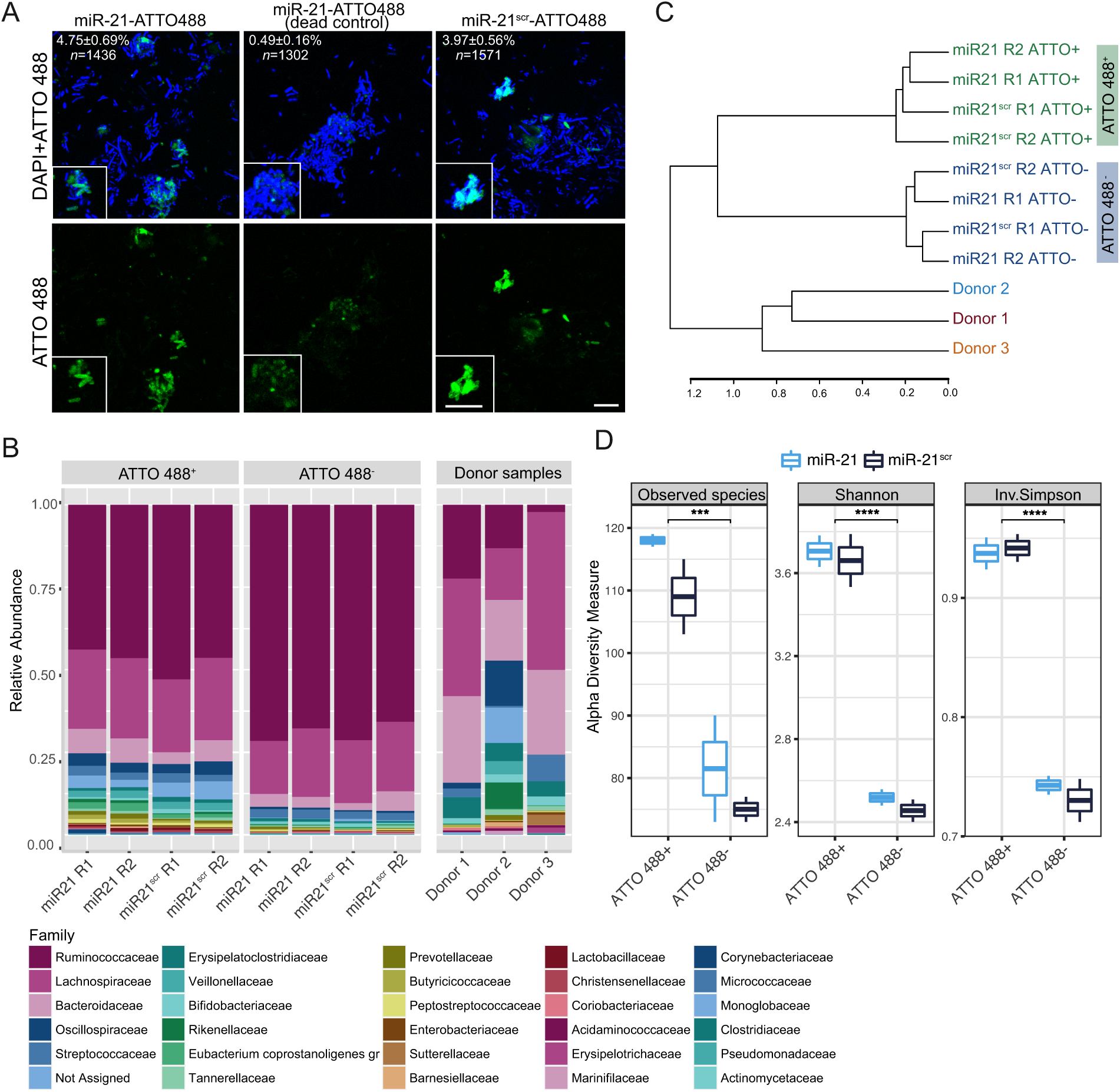
Imaging, sorting and sequencing of microbial populations associating or not with fluorescently labelled miRNAs. (**A**) Fluorescence microscopy imaging of gut microbiota cells labelled with the nucleic acid dye DAPI (shown in blue) after one hour of incubation with miR-21- or miR-21^scr^-ATTO 488 (in green). Representative images of fixed microbiota cells (dead control) incubated with miR-21-ATTO 488 are shown in the middle panel. Numbers in the top left corner indicate the percentage of cells displaying green fluorescence and *n* indicates the total number of cells analyzed. Scale bar: 10 µm. (**B**) Family-level relative abundance profile of communities present in ATTO 488^+^ or ATTO 488^-^ sorted fractions, obtained after incubation with miR-21- or miR-21^scr^-ATTO 488. Incubations were established from a mix of faecal samples originating from three donors, whose individual faecal microbiome composition is shown in the “Donor sample” panel. Each bar represents the mean from two replicates. (**C**) Dendrogram summarizing the hierarchical clustering of microbial communities present in ATTO 488+ or ATTO 488- sorted fractions, as determined by 16S rRNA gene amplicon sequencing. Microbial communities present in fractions sorted based on the ATTO 488 positive (ATTO 488^+^) gate are highlighted in green, while the ones based on the ATTO 488 negative gate (ATTO 488^-^) are highlighted in blue. Communities of the three individual donors from which a combined faecal slurry community was obtained are also shown as a reference. Data from two replicates per miR and per gate are shown: R1 and R2. (**D**) Alpha diversity metrics (Observed ASVs, Shannon index and Inverse Simpson’s diversity index) in gut microbial communities described in (B) and (C). Boxes represent the median, first and third quartile. Whiskers extend to the highest and lowest values that are within one and a half times the interquartile range. ***p<0.001, ****p<0.0001; paired Wilcoxon test.

### A large and diverse subset of the faecal microbiota interacts with miRNAs

To determine which microbial taxa associate with miR-21 and elucidate if these taxa differ from taxa interacting with miR-21^scr^, we performed faecal microbiota incubations with fluorescently tagged versions of miR-21 or miR-21^scr^. Fluorescence microscopy analysis of samples incubated with ATTO488-labelled miRNAs revealed a strong fluorescence signal originating from both miR-21 or miR-21^scr^ that colocalized with the nucleic acid stain DAPI in a fraction of microbial cells (**Fig. 2A**). Fluorescent signal originating from miR-21 incubations with dead microbial cells yielded a much less intense and scattered signal that did not show a strong colocalization with nucleic acids (**Fig. 2A**, middle column). To determine the phylogenetic identity of fluorescently labelled cells, samples were subjected to fluorescence-activated cell sorting (**Fig. S3**). We sorted a similar number (25,000) of cell events within ATTO488^+^ (interacting with fluorescently labelled miRNAs) and ATTO488^-^ (not interacting with miRNAs) gates. Sorted fractions were subsequently processed for 16S rRNA gene amplicon sequencing to identify microbial populations within each fraction (**Tables S3 and S4**). This analysis revealed that both sorted fractions were dominated by microbial taxa belonging to the families *Ruminococcaceae*, followed by *Lachnospiraceae* and *Bacteroidaceae* (**Fig. 2B**). These families were also dominant in faecal sample material that was used to establish these incubations (**Fig. 2B**, “Donor samples”, **Table S3**). Hierarchical clustering of microbial community composition at the ASV level revealed that ATTO488^+^ fractions cluster together, forming a separate cluster from ATTO488^-^ fractions (**Fig. 2C**). This is an indication of the presence of a unique microbial subpopulation which is capable of miRNA interaction. However, the miRNA sequence itself does not appear to be a significant driver of the taxonomic composition of sorted cells (**Fig. 2C**). Alpha diversity analyses revealed that both miR-21- and miR-21^scr^- interacting (ATTO488^+^) fractions are significantly more diverse and richer than non-interacting fractions (ATTO488^-^) (**Fig. 2B,D**). This suggests a larger fraction of the faecal microbiota associates with fluorescently-labelled miRNAs than does not, and that this occurs in a miRNA sequence-independent manner.

To gain further insights into the microbial taxa interacting with miR-21 and miR-21^scr^, we performed differential abundance analysis using DESeq2 (42), to retrieve all taxa significantly enriched in ATTO488^+^ sorted fractions, independently of the sequence of the miRNA supplemented. ASVs belonging to the genera *Limosilactobacillus*, *Ruminococcus*, *Coprococcus* or *Clostridia UCG-14* were among the most significantly enriched (Log2FC>1 and padj<0.05) across all ATTO488^+^ sorted fractions (**Fig. 3A, Table S5**). Genera such as *Collinsella, Erysipelotrichaceae UCG-003, Pseudomonas* and *Faecalibacterium*, on the other hand, are significantly enriched in ATTO488^-^ fractions (Log2FC<-1 and padj<0.05) indicating that these taxa interact to a lower extent, or not at all, with miR-21 or miR-21^scr^ (**Fig. 3A**). The genera *Streptococcus*, *Bacteroides* and *Lachnospiraceae_unclassified* were found to have variable trends, with some ASVs being enriched while others were depleted in ATTO488^+^ fractions (**Fig. 3A**).

**Figure 3.**
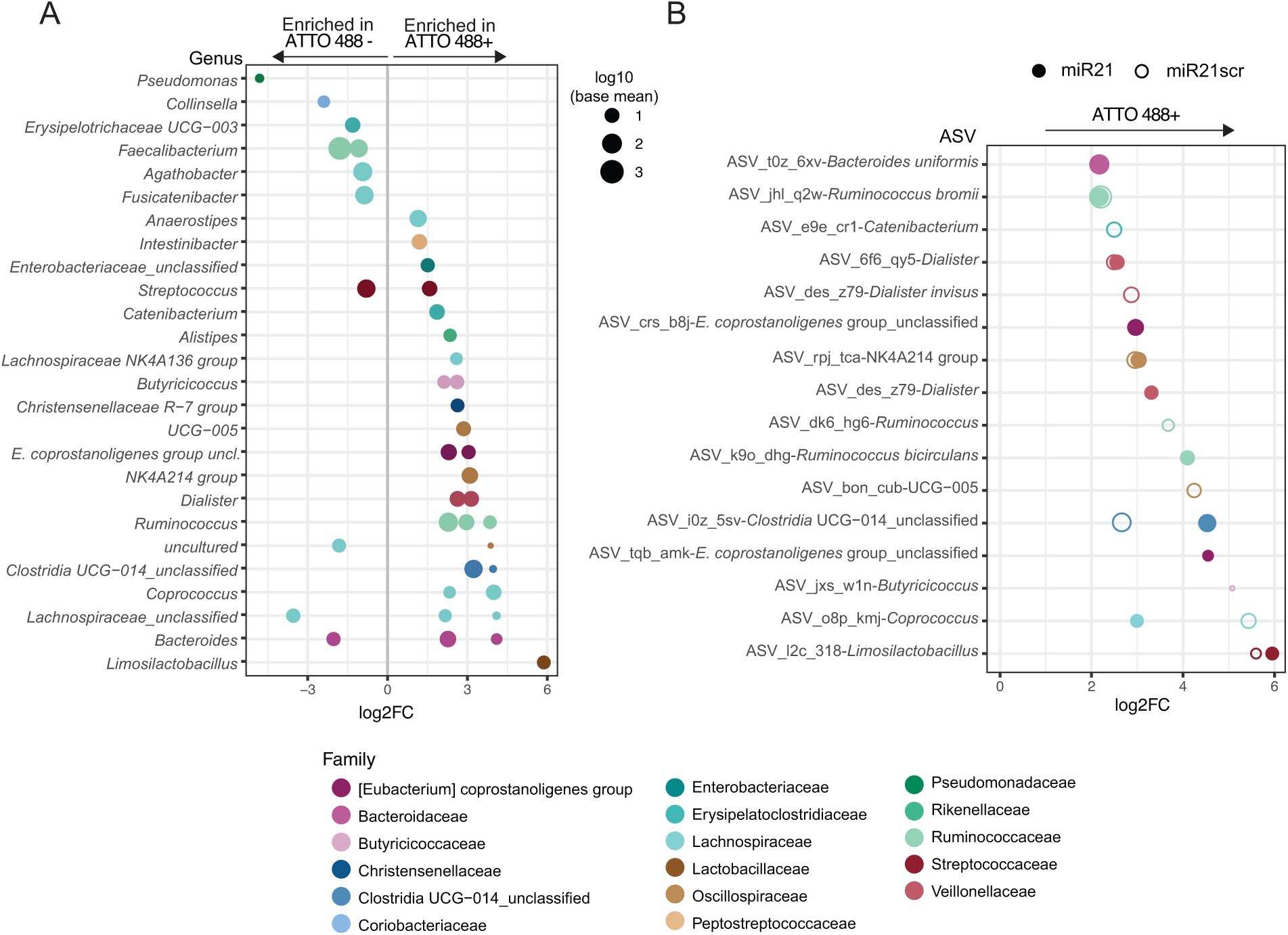
Microbial taxa associating with fluorescently-labelled miR-21 and miR-21^scr^. (**A**) DESeq2-based differential abundance plot showing all ASVs whose relative abundance significantly differs across both miR-21 and miR-21^scr^ ATTO 488^+^ gated sorts versus miR-21 and miR-21^scr^ ATTO 488^-^ gated sorts. Each data point represents an ASV grouped by genus and coloured by family. (**B**) DESeq2-based differential abundance analysis of bacterial populations of sorted fractions for either miR-21 or miR-21^scr^. All ASVs whose abundance is significantly higher in miR-21-ATTO 488^+^ compared to miR-21-ATTO 488^-^ sorts (filled circle), or in miR-21^scr^-ATTO 488^+^ sorts compared to miR-21^scr^-ATTO 488^-^. In (A) and (B), the size of each data point represents the base mean counts for that ASV across the dataset (DeSeq2 analyses). The x-axis represents the log2 fold change in abundance of the respective ASV across treatments. Only ASVs with an adjusted p-value <0.05 are shown.

To better understand any preferences of microbiota in interacting with a particular miRNA sequence, we also performed differential abundance analysis comparing miR- 21-ATTO488^+^ versus miR-21-ATTO488^-^ fractions, and miR-21^scr^-ATTO488^+^ versus - miR-21^scr^ ATTO488^-^ fractions (**Fig. 3B, Table S5**). A total of 11 ASVs were found to be significantly enriched (Log2FC>1 and padj<0.05) in ATTO488^+^ fractions incubated with fluorescently labelled miR-21 (**Fig. 3B**, closed circles). Also, 11 ASVs were found significantly enriched in ATTO488^+^ fractions incubated with fluorescently labelled miR- 21^scr^ (**Fig. 3B**, open circles). Six of these ASVs were found to be enriched in both miR- 21- and miR-21^scr^-ATTO488^+^ fractions (**Fig. 3B**, ASVs displaying both open and closed circles). These belong to the genera *Limosilactobacillus*, *Coprococcus*, *Clostridia* UCG- 014, *Lachnospiraceae* NK4A214 group, *Dialister* and *Ruminococcus.* ASVs significantly enriched in miR-21-interacting fractions only belong to the genera *Ruminococcus*, *Eubacterium coprostanoligenes group_unclassified*, *Dialister* and *Bacteroides uniformis* (**Fig. 3B**). This suggests that there is a large overlap between taxa interacting with miR-21 and miR-21^scr^, but that a narrow group of microbial taxa may preferentially interact with miR-21.

### miR-21 specifically alters messenger RNA levels in *B. thetaiotaomicron*

MiR-21 depletion has been previously shown to alter gut microbiota composition and ameliorate DSS-induced colitis in mice (43). Our results suggest that taxa implicated in intestinal pathogenesis including *Bacteroides* and *Ruminococcus* may preferentially interact with miR-21. We therefore investigated whether miR-21 has the potential to alter the transcription of genes in these members of the microbiota. Thus, we performed transcriptomic analysis of a prominent gut *Bacteroides* species - *B. thetaiotaomicron* - supplemented with miR-21. In addition to a miR-21^scr^ control, a double-stranded small RNA oligonucleotide (see Methods) was also included as a control, to determine if there is a specific effect of miRNA mimics compared to small RNA supplementation. Supplementation with miR-21, miR-21^scr^ and control small RNA molecules did not significantly impact *B. thetaiotaomicron* growth compared to the water control (**Fig. 4A**). Depletion of supplemented miRNAs from supernatant fractions was rapid, with qPCR results indicating that only 0.11% of the initial miR concentration remained in the vials one hour after supplementation with either miR-21 or miR-21^scr^ (**Table S6**). Principal component analysis of *B. thetaiotaomicron* transcriptomic profiles of samples collected one hour after miRNA or small RNA supplementation revealed a significant impact of treatment on *B. thetaiotaomicron* transcription (**Fig. 4B**; p<0.001, PERMANOVA). The water control and small RNA supplementation samples clustered closer together and away from either miRNA supplemented samples, indicating that the transcriptomic profile induced by miRNA supplementation is distinct from that induced by the small RNA control (**Fig. 4B**). Differential expression analysis revealed that a total of 185 transcripts were significantly differentially abundant (log2 fold change <-1 or >1, adjusted p-value<0.05, DESeq2 analyses) in miR-21 incubation compared to the water control (**Fig. 4C**). Supplementation with miR-21^scr^ resulted in 63 differentially abundant transcripts (**Fig. 4D**), whereas supplementation with the small RNA control significantly altered the levels of only 3 transcripts compared to the water control (**Fig. 4E**). Of the 185 transcripts differentially abundant in the presence of miR-21, 10 were significantly increased, while 175 showed a significant decrease in abundance (**Fig. 5A**). A significant fraction of the latter (49) is also differentially less abundant upon miR-21^scr^ supplementation (**Table S7**). These 49 transcripts potentially represent the overall impact that supplementation of miRNA mimics (independent of the sequence) has on *B. thetaiotaomicron* transcript abundance profile (**Fig. 5A**). While several of these miRNA-regulated genes are uncharacterized, many of them encode for DNA binding proteins, ribosomal proteins, transcriptional regulators and transfer RNAs (tRNAs) (**Table S6**). Transcripts whose abundance was specifically decreased upon miR-21 supplementation comprise a putative ferric citrate regulator (BT_4356), a putative rubredoxin (BT_2539), a transcriptional regulator from the Mar family (BT_2435) and genes involved in the regulation of iron dicitrate transport (BT_2462). The three transcripts with decreased abundance upon small RNA supplementation are part of a putative cation efflux system (BT_4693-BT_4695) (**Table S7**).

**Figure 4.**
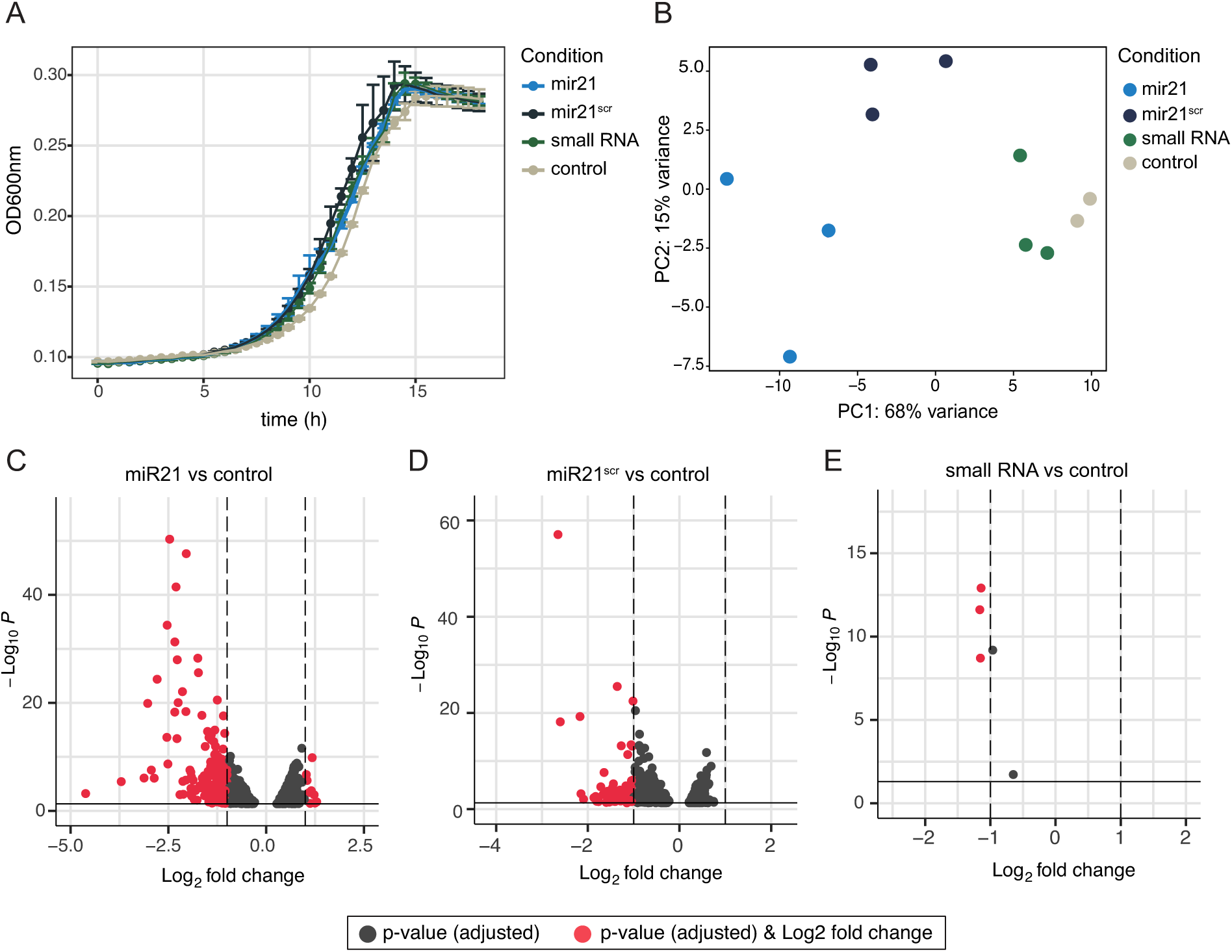
Transcriptomic analyses of *B. thetaiotaomicron* cells supplemented with miR- 21 or controls. (A) Growth of *B. thetaiotaomicron* in the presence of 250 nM of miR-21, miR- 21^scr^, a small RNA control and no amendment (water) control. Data from 3 independent growths per condition is shown. Dots depict the mean per condition, and error bars represent the standard deviation. (**B**) Principal coordinate analysis (PCoA) based on Bray-Curtis distances summarizing transcript level profiles (read counts) of *B. thetaiotaomicron* cells collected one hour after amendment with each indicated small RNA (or water as a control). (**C-E**) Volcano plots representing differentially abundant *B. thetaiotaomicron* transcripts incubated with miR- 21 (**C**), miR-21^scr^ (**D**), and small RNA control (**E**) compared to the water control. In all volcano plots, the x-axis represents the log2 fold change and the y-axis represents -log10 (adjusted p-value). All transcripts were identified and analyzed using the DESeq2, and transcripts with an adjusted p-value < 0.05 are represented (grey). Transcripts that show both a log2 fold change > 1 or < -1 and have a p-value < 0.05 are shown in red.

**Figure 5.**
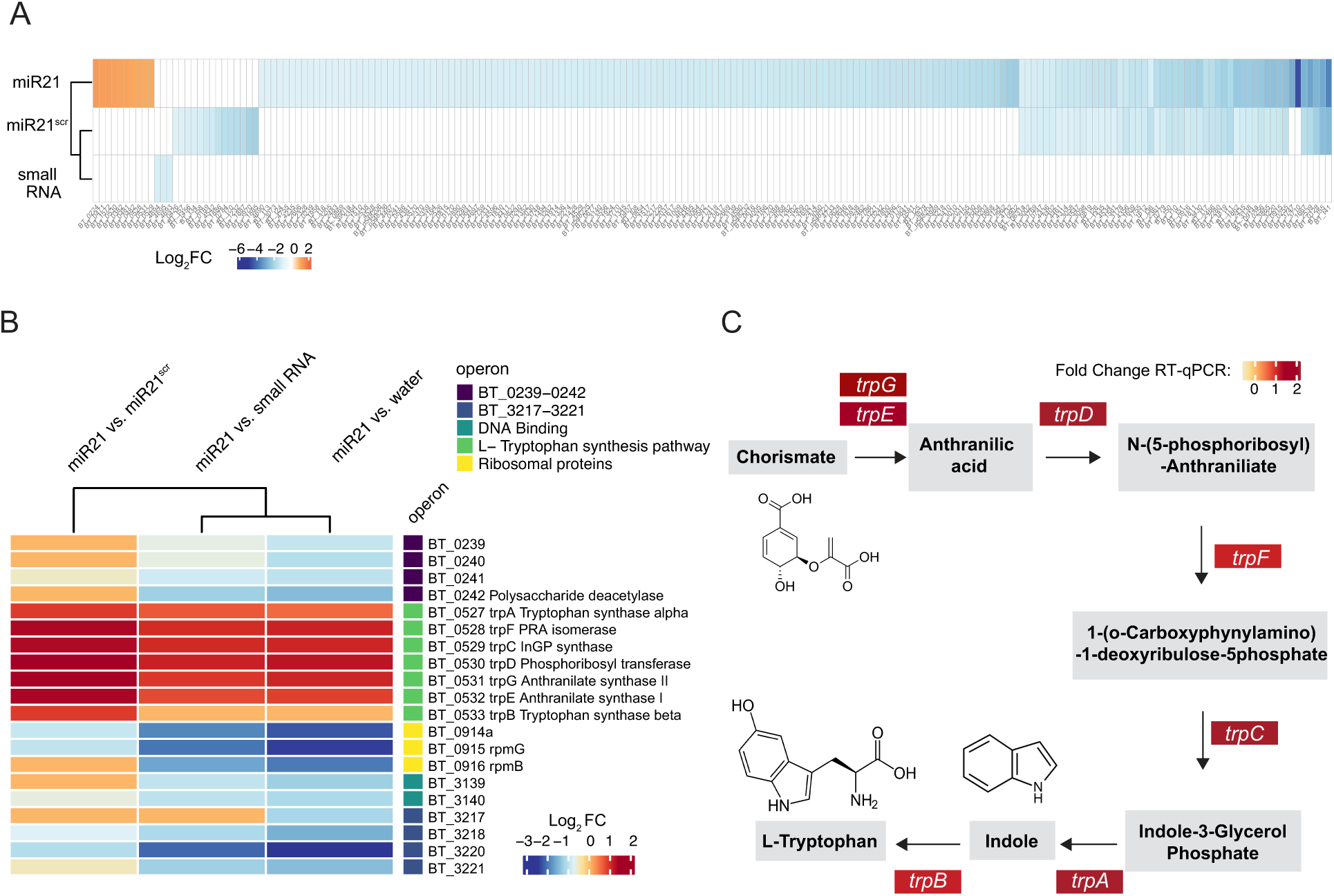
MiR-21 differentially impacts *B. thetaiotaomicron* transcription and regulates transcript levels of a gene cluster involved in indole and L-tryptophan synthesis. (**A**) Heatmap showing significantly differentially expressed genes (DEGs) (adjusted p-value <0.05) between each indicated small-RNA amended sample (relative to the water control). Only genes with a log2 fold change > 1 or < -1 are shown. (**B**) Heatmap summarizing the log2 fold change in transcript levels in cells supplemented with miR-21 compared to all other treatments. (**C**) Representation of the enzymatic steps involved in indole and L-tryptophan synthesis from chorismate highlighting intermediates and genes encoding enzymes catabolizing each step in *Bacteroides* spp (44). Colour boxes indicate the fold change in mRNA levels relative to the water controls, as determined by RT-qPCR.

Transcripts significantly upregulated upon miR-21 supplementation are part of operon BT_0527-BT_0533 (**Fig. 5B**), which encodes seven genes necessary for the biosynthesis of indole and L-tryptophan from chorismate and L-glutamine (*trp*AFCDGEB) (44) (**Fig. 5C**). Furthermore, miR-21 was also shown to boost levels of these transcripts in comparison to the addition of miR-21^scr^ or the small RNA control (**Fig. 5B**), highlighting the specific impact that miR-21 supplementation has on this pathway. The increased expression of *trp*AFCDGEB upon miR-21 supplementation was confirmed by RT-qPCR (**Fig. 5C, Table S8**). Transcripts encoded by four additional gene clusters, *i.e*., genes present sequentially on the same strand, were found to be decreased in the presence of miR-21 compared to the water control (**Fig. 5B**). One of these clusters is part of the ribosomal operon *rpmB-rpmG* and encodes the ribosomal proteins L33 and L28 (BT_0914a-BT_0916). Additional gene clusters encode proteins with motifs involved in DNA binding and transcriptional regulation (BT_3139-BT-3140) and virulence (BT0239-BT0242). However, the specific function of most of the proteins encoded by these genes is unknown outside of their sequence motifs (**Table S7**). None of these clusters was identified as consistently differentially expressed upon miR-21 supplementation compared to all other conditions. Collectively, these results demonstrate a selective increase in tryptophan and indole biosynthesis transcripts in *B. thetaiotaomicron* by miR-21.

## Discussion

There is growing evidence linking faecal miRNAs and the gut microbiota to gastrointestinal diseases, but less research has been done to determine whether and how miRNAs and the gut microbiota interact to influence gut function. Here we investigated how the host can influence microbiome function through miR-21. Our time-resolved analysis of the dynamics of miR21-microbiota interactions demonstrated the ability of miR-21 and a scrambled control to associate with diverse microbial cells within complex gut communities in a rapid and sequence-independent manner (**Fig. 1 and 2**). Most miRNAs found in the gut lumen are thought to be present inside vesicles secreted by intestinal epithelial cells and may enter microbial cells following the fusion of these vesicles with the bacterial membrane through a process that is yet to be described (45). However, in this and other (35, 41) studies, miRNAs were supplemented in a naked state, i.e., were not enclosed in vesicles. How naked miRNAs enter microbial cells is still unknown, but their entry may be connected to bacterial natural competence (46).

More research is needed to fully understand the processes governing the entry of miRNAs into bacteria, although it is reasonable to anticipate that these mechanisms will be ubiquitous, and incapable of distinguishing miRNAs based on their sequence, in agreement with our results. The ability to uptake miRNAs is also expected to be common to several bacterial taxa, and our results demonstrate that miRNAs interact with and are potentially internalized by a large fraction of the gut microbiome (**Fig. 2D**). Nevertheless, miR-21 appears to maintain a preferential association with a few taxa including *Dialister*, *Bacteroides, Eubacterium,* and *Ruminococcus* species. Interestingly, a miR-21 deletion mutant mouse exhibits a shifted microbiome with lower levels of *Bacteroides* and *Ruminococcus* (43). Others have shown that oral administration of miR-30d to mice regulates the expression of a lactase enzyme in *Akkermansia muciniphila* and results in an increased abundance of *A. muciniphila* in the gut (37). More research is needed to understand if the preferential association of *Bacteroides* and other taxa with miR-21 can shape their abundance and/or fitness in the gut environment.

This study further demonstrates that miR-21 supplementation results in altered levels of 185 transcripts in the gut symbiont *B. thetaiotaomicron*, the majority of which were present in lower abundances compared to controls. These results support several other studies that suggest that miRNAs control gene expression through the degradation of bacterial mRNA (47). It is not understood how exactly miRNAs regulate transcript levels in bacteria, but it is possible that miRNAs act similarly to *trans* encoded bacterial small RNAs, which can form base pairing with multiple target mRNAs and result in a global regulation of a physiological response (48). Of note, supplementation with either a miR-21 scramble, or a small RNA control did not induce a comparable magnitude of differences in gene transcript levels as observed for miR-21 (**Fig. 5**). Thus, miR-21 has the capacity to precisely modify transcription in this gut commensal, strongly pointing to the existence of a miR-21-microbiota axis.

Notably, levels of *trpAFCDGEB* transcripts encoding various enzymes involved in indole and L-tryptophan (Trp) biosynthesis (44) were shown to be increased in miR-21- supplemented *B. thetaiotaomicron* (**Fig. 5**). In *E. coli*, these genes are co-transcribed under the control of the tryptophan repressor *trpR* (44). It is plausible that miR-21 targets a yet-to-be-identified homologue of such transcriptional repressor in *B. thetaiotaomicron,* resulting in the observed increase in *trp*AFCDGEB transcript levels (49). Tryptophan is an essential amino acid that cannot be synthesized *de novo* by human cells and instead is acquired via the diet. Intestinal tryptophan metabolism is under direct or indirect control of the microbiota and results in the production of various catabolites that regulate key gastrointestinal functions such as intestinal immune homeostasis, permeability, and motility (50, 51). The majority of the body’s serotonin is synthesized from tryptophan in the gut by enterochromaffin cells (EC), and serotonin release by ECs activates peristalsis and secretion (51, 52). Changes in tryptophan metabolism leading to increased serotonin production were found to be positively correlated with gastrointestinal symptoms in patients with IBS (38). Notably, administration of a bacterial strain enzymatically capable of synthesizing tryptophan promotes intestinal motility (53). Our findings imply that miR-21 may control tryptophan synthesis and gastrointestinal motility by modulating *trp*AFCDGEB expression in *B. thetaiotaomicron*, and perhaps also in other microbiome members.

Most reports on the dysregulation of miRNAs in IBS have been conducted in tissue and cell cultures, and there are currently few, if any, studies on the quantitative assessment of stool miRNAs in IBS patients (54). Here we show that lower and variable levels of miR-21 were detected in the stool of a small cohort of IBS patients compared to controls (**Fig. 1A**). These results may help to explain the variability in gastrointestinal motility experienced by IBS patients and open the door for future efforts to explore the role of other miRNAs in mediating microbe-host interactions.

Tryptophan catabolites, including indole and derivatives such as indoleacrylic acid, are also known to enhance intestinal epithelial barrier function and attenuate inflammatory processes in mice by promoting goblet cell differentiation and mucus production, in part through the activation of the aryl hydrocarbon receptor (51, 55). Patients suffering from IBD have reduced serum tryptophan concentrations (56), and a tryptophan-free diet increases susceptibility to DSS-induced inflammation in mice (57). The elevated levels of miR-21 in the stool of IBD patients (30, 31) may reflect a host response to counteract tryptophan and indole deficits by promoting their production by the microbiota. Thus, miR-21 may regulate gut inflammation also indirectly, via the microbiome. Overall, our results highlight the importance of considering miRNA-microbiota interactions in the context of gut health and disease, and identify microbial members and molecules that could be the target of future interventions for gut disorders.

## Materials and Methods

### Faecal sample collection

Faecal samples from patients undergoing colorectal cancer screening colonoscopy at the Vienna General Hospital were collected from the first stool after starting polyethylene glycol-based bowel preparation. Samples were stored at 4 °C overnight and transferred to -80°C upon arrival at the hospital. This study was conducted under the Health Research Authority of the Ethics Committee of the Medical University of Vienna, ethics approval number 1617/2014. All study participants gave written informed consent before study inclusion. The study was conducted following the ethical principles of the Declaration of Helsinki.

Fresh faecal samples for miRNA supplementation incubations were collected from three adult individuals with no history of inflammatory disease and who had not taken antibiotics in the previous 3 months. All participants signed an informed consent form and self-sampled using a sterile polypropylene tube with the attached sampling spoon (Sarstedt, Nümbrecht, DE). Samples were transferred into an anaerobic chamber and further processed within 90 minutes of defecation. The study protocol was approved by the University of Vienna Ethics Committee (reference #00161). All data is completely anonymized and adheres to the University regulations.

### Total RNA isolation from faecal samples

For extraction of total nucleic acids (TNA) from patient stool samples, ∽200 mg of frozen material was weighed directly into Lysing Matrix E 2 ml screw-cap tubes (MP Biomedicals) containing 500 μl of centrimonium bromide (CTAB) extraction buffer (0.35 M of NaCl, 0.12 M of CTAB, 0.008 M of KH2PO4, 0.17 M of K2HPO4 ·3H2O). TNA were extracted using the phenol:chloroform method. Samples were subjected to DNAse treatment with TURBO DNAse I (Invitrogen), according to the manufacturer’s instructions. RNA samples treated with DNAse I were subjected to purification using the RNA Clean and Concentrator^TM^ kit (Zymo research) following the protocol provided by the manufacturer for purification and concentration of total RNA including small RNAs. All samples were eluted in 15 μl of nuclease-free water and stored at -80°C. RNA samples were quantified using the Invitrogen™ Qubit™ 4 Fluorometer and Qubit™ HS RNA Assay kit (Thermo Fisher Scientific), according to the manufacturer’s instructions.

### Quantification of native miRNAs in faecal sample material

Polyadenylation was performed before reverse transcribing miRNAs into cDNA by mixing 1x of PolyA polymerase buffer, 0.5 mM of rATP, 1 U of PolyA polymerase (New England Biolabs GmbH) and 1.5 μg of RNA, according to the manufacturer’s instructions. A murine leukaemia virus (MuVL) transcriptase (New England Biolabs GmbH) was used to transcribe polyadenylated RNA into cDNA using the reverse transcription adaptor primer 5’- CAGGTCCAGTTTTTTTTTTTTTTTVN -3’ (58). Reactions were prepared by mixing 1x of RT buffer, 10 mM of dNTP mix, 0.5 μM of dT adaptor primer, 2 U μL^-1^ of RNAse OUT, 10 U μL^-1^ of M-MuVL reverse transcriptase. cDNA synthesis was performed in a thermal cycler (T100, BioRad). All reactions were further diluted by adding 130 μl of nuclease-free water and stored at -20°C.

For quantitative PCR targeting the miRNAs hsa-miR-21-5p and hsa-miR-26b-5p, a master mix was prepared by mixing 1x iQ SyBr Green mix (BioRad), 0.4 μM of primer forward and reverse (**Table S2**), 0.4 μM of BSA, 2 μl of the template cDNA and nuclease-free water to bring reaction volume up to 10 μl. Samples were placed into a Real-Time PCR detection system CFX96 (BioRad) and incubated at 95°C for 5 minutes, followed by 40 cycles of 95°C for 15 sec, 60°C for 45 sec and a plate read. Each sample was run in triplicates. An aliquot of each cDNA sample was pooled into a new tube to produce a mixture of cDNAs from all samples. This mixture was serially diluted to obtain 10^-1^, 10^-2^, 10^-3^, 10^-4^ and 10^-5^ dilutions of cDNAs and used to determine qPCR amplification efficiencies for each miRNA primer set. Amplification efficiency was calculated for each set of standards, and only reactions with efficiencies between 80-110% were considered (58).

### Preparation of faecal slurries and dead control

To establish incubations with miRNAs, faecal samples were freshly collected and introduced in a Coy anaerobic chamber (85% N2, 10% CO2, 5% H2, Coy Labs, USA) within 90 minutes of defecation. All other reagents and laboratory equipment to process samples were introduced in the anaerobic chamber one day before experimentation to reduce oxygen levels. A 10% (w/v) faecal slurry was prepared by adding 1 x phosphate-buffered saline (PBS) to the sample, followed by thorough vortexing for 2 minutes. The sample was then left for 10 minutes, so that larger particles could settle to the bottom of the vial. 1 mL of the pre-settled mixture was transferred to a new 50 mL falcon tube and diluted ten times with sterile M9 mineral medium (prepared with nuclease-free water) supplemented with 0.5 mg.mL-1 D-glucose (Merck), 0.1% L-cysteine, 0.5% v/v of vitamin solution and 0.1% (v/v) mineral solution (both prepared according to DSMZ, Medium 461), referred to as supplemented M9 (sM9). The mixture was thoroughly vortexed, and again larger particles were allowed to settle. Aliquots of this faecal slurry were used for subsequent experiments. For EtOH fixation (dead microbiota control), an aliquot of the sample was incubated in 50% (v/v) ethanol at -20°C for 1 hour. The mixture was then spun down for 5 minutes at 4°C, 12000 *g*, the supernatant was discarded, and the recovered pellet was reintroduced in the anaerobic chamber, allowed to re-equilibrate for 1 hour, and finally resuspended in the same initial volume of anaerobic sM9 medium.

### Faecal sample incubations with miRNA mimics

All tubes and pipette tips used to set up incubations were nuclease-free, to prevent miRNA degradation. Human miR-21 (hsa-miR-21-5p, Sequence: 5’ – UAGCUUAUCAGACUGAUGUUGA - 3’; HMI0371 MISSION® microRNA Mimic, Merck), or miR-21 scramble control (Sequence: 5’ - GCAUAUUCGGUCGUUAUAAGAU - 3’; custom designed MISSION® microRNA Mimic, Merck) were diluted in water, and added to the faecal slurries to a final concentration of 250 nM. Data on the absolute concentration of miRNAs in stool are scarce. We found data reporting the presence of 20 pmol of miR-30d in a faecal pellet (25 mg) of a mouse model of multiple sclerosis (41). The authors also reported that miR-30d levels in diseased mice are approximately twice as in healthy control mice. Thus, assuming 10 pmol miR-30d are present in a 25 mg pellet from a healthy mouse and a pellet density of 1.06 g.mL^-1^ (59), the estimated concentration of miR-30d in mouse faeces is approximately 425 nM. We estimated that miR-21 is present in faeces in the high nanomolar range and therefore tested supplementation at a concentration of 250 nM. A water control where the same volume of nuclease-free water only (same water used to dilute the miRNA mimics) was also included, and incubations were set in duplicates. A sample was collected immediately after (T0) or at 1, 2, 4, and 6 hours of incubation. Samples were spun down for 5 minutes (4°C at 12000 *g*), and the supernatant was removed and stored at -80°C (**Fig. 1B-F**). The cell fraction (pellet) was washed with one volume ice-cold 1 x PBS and spun down again for 5 minutes (4°C at 12000 *g*). The supernatant was discarded, and the pellet was stored at -80°C.

To determine if the depletion of miRNA over time in pellet fractions was due to RNase activity, live faecal slurry incubations with miR-21 were set up with or without amendment of 2 U.µl^-1^ of an RNAse inhibitor (RNaseOUT™, Thermo Fisher Scientific), and sampled as described above. To determine the stability of miR-21 in the incubation medium, miR-21 (250 nM) was added to vials containing M9 medium only (no microbiota), and supernatants were collected as described above and stored at -75°C for further analysis.

### Total RNA isolation from faecal sample incubations

The Total RNA Purification Kit (Norgen Biotek Corp) was used to isolate and purify total RNA from the supernatant and cell fractions, including miRNAs. To improve cell lysis in such a complex microbial sample, cell pellet fractions were resuspended in 100 μL of TE-Lyso lysis buffer containing: 20 mM Tris-HCl buffer pH = 8.0, 2 mM EDTA, 1.2% Triton-X, nuclease-free water and 20 mg.mL^-1^ of lysozyme from chicken egg, freshly added (Merck). Lysis was carried at 37°C for 30 minutes. Following this step, pellet and supernatant fractions (both 100 μL) were treated the same for all downstream steps, and total RNA was extracted following the manufacturer’s instructions for total RNA isolation. RNA was finally eluted in 25 μL RNase-free water and stored at -80°C.

### Quantification of microRNA mimics

TaqMan™ MicroRNA assays (Applied Biosystems™) were used to quantify miR-21 (assay ID 000397) and miR-21^scr^ (assay ID CT7DPW3, custom-designed) in the supernatant and pellet fractions. These assays involve reverse transcription (RT) with miRNA-specific primers, followed by real-time PCR with TaqMan probes. For reverse transcription, RNA samples were diluted 1:10 in nuclease-free water and 5 μL of the diluted RNA product was mixed thoroughly with 3 μL of the 5 X RT primers and RT was carried according to the manufacturer’s instructions using a TaqMan™ MicroRNA Reverse Transcription Kit (Thermo Fisher Scientific) in a thermal cycler (Bio-Rad Laboratories). As a control, a mock cDNA synthesis reaction was also performed, in which no reverse transcriptase was added to the samples of interest. Following completion of the reactions, the cDNA products were stored at -20°C or stored on ice and immediately used for quantitative PCR.

To precisely quantify miRNAs in faecal incubations, qPCR was performed using miR-specific probes provided by the TaqMan™ MicroRNA assays (catalogues numbers indicated above) and a TaqMan™ Fast Advanced Master Mix (Thermo Fisher Scientific), according to the manufacturer’s instructions. In addition to standards, a negative control using only nuclease-free water was included to ensure no contamination of the master mix. All samples and controls were run in triplicates in a CFX96™ real-time PCR system (Bio-Rad Laboratories). Standard curves were produced by generating a 10-fold serial dilution of each miRNA mimic, starting with a sample at a concentration of 250 nM. RNA extraction, reverse transcription and qPCR reactions were set up as described above. Standard curves were then generated using the results from the qPCR and mapping miRNA concentration of the standards against Cq value. The acquired regression line equation was then used to calculate miRNA concentration in all the samples based on their respective Cq values.

### Faecal sample incubations with fluorescently labelled miRNAs

Three different faecal samples were collected from the same three donors used in the previous miRNA incubation experiments. Faecal sample collection and faecal slurry preparations were carried out as described above. An aliquot (0.5 mL) of each slurry was pelleted by centrifugation, and immediately stored at -80°C for downstream DNA extraction to profile the microbiome of each donor. Following this step, the faecal slurries from all 3 donors were combined in equal volumes in a 50 mL falcon tube, to create a single faecal slurry representative of all 3 donors for the experiment. The faecal slurries were aliquoted into tubes and 250 nM of ATTO 488-tagged Mission MicroRNA mimics (Sequence: 5’-[ATTO488]UCAACAUCAGUCUGAUAAGUCUA [dT][dT]-3’) and miR-21^scr^ (Sequence: 5’-[ATTO488]AUCUUAUAACGACCGAAUAUUGC[dT][dT]-3’; both from Merck) were added. Negative control incubations were set by adding the same volume of nuclease-free water to each tube. All samples (water control, miR-21, miR-21^scr^) were set up in duplicates. The samples were gently vortexed and incubated at 37 °C for 30 minutes. After the incubation, the samples were vortexed, washed with 1 x PBS and the pellet was re-suspended in 1 mL 1 x PBS. The samples were put on ice and removed from the tent for ethanol fixation as described above. For the miRNA- amended samples, each sample replicate was set up and processed one at a time to prevent miRNA degradation by the bacterial cells and minimize fluorescent dye bleaching. During all steps of the workflow, all samples were carefully protected from light to preserve the fluorescence signals.

### Fluorescence microscopy

Ethanol-fixed samples (10 μL) were spotted onto a glass slide and let to dry at 46°C. Samples were DAPI stained for 10 minutes using 20 μl of 1 μg/ml DAPI (Sigma-Aldrich) per well before washing once in ice-cold MiliQ water. Slides were subsequently dried with pressurized air and a drop of CitiFluor™ AF1 anti-fading agent was added before applying the coverslip. Images were acquired on a confocal TCS SP8X microscope (Leica Microsystems, Germany), using a 63x glycerol objective. Images were analyzed using ImageJ.

### Fluorescence-activated cell Sorting (FACS)

Ethanol-fixed samples incubated with ATTO 488-labelled miRs were stained by adding 2 μM SYTO™ 62 red fluorescent nucleic acid stain (Thermo Fisher Scientific) followed by incubation in the dark for 30 minutes. Samples were analyzed on a BD FACSMelody^TM^ Cell Sorter, calibrated on the day according to the manufacturer’s instructions. Following system calibration, all samples were transferred into flow cytometry tubes and passed through a 35 µm pore cell strainer cap (Corning), to remove larger particles. At times when the concentration of cells was too high, the samples were diluted by adding one volume of 1x PBS. Background signals from the instrument and PBS were identified using the operational parameters forward scatter (FSC) and side scatter (SSC). Singlet discrimination was performed. Gating for microbial cells was based on the presence of SYTO™ 62 signal, established based on the PerCP-Cy5.5 fluorescence-forward scatter plot, by comparing SYTO™ 62 stained and unstained samples. The gates for the ATTO 488^+^ and ATTO 488^-^ populations within SYTO™ 62^+^ events were established based on the Alexa488 fluorescence- forward-scatter plot by directly comparing the distribution of events from samples incubated with water with samples incubated with ATTO 488-miRs (**Fig.S3**). Fluorescence signals were detected using the blue 488 nm optical laser and a 527/32 (ATTO 488) or a 665 LP filter (SYTO™ 62). Gated events were sorted on purity mode. Instrument parameters and gating strategy were kept constant during the acquisition of all samples. All sorts, as well as two aliquots of the 1 x PBS, used to dilute the samples were subsequently stored at -80 °C for downstream processing.

### DNA isolation of FACS cells and 16S rRNA gene amplification and sequencing

For DNA isolation from sorted cell fractions, samples were defrosted on ice and 180 μL of TE-Lyso lysis buffer (as described above) was added. Samples were incubated for 45 min at 37°C with gentle shaking (400 rpm) in a thermal block. Before extraction, all kit buffers and TE lysis buffer (before lysozyme addition) were UV-radiated for 30 minutes in a PCR Workstation. The DNA was then extracted using the innuPREP DNA Mini Kit (Analytik Jena GmbH) according to the manufacturer’s instructions. Blank DNA extractions (buffer only) as well as extractions from 2 aliquots of 1xPBS used in FACS were performed in parallel. Pelleted faecal slurries from the 3 individual donors used to set up miRNA incubations were also extracted using the same protocol.

Amplification of bacterial and archaeal 16S rRNA genes from DNA samples extracted from sorted cells, faecal slurries and controls was performed with a two-step barcoding approach (60) using V4 primers 515F (5’-GTGYCAGCMGCCGCGGTAA-3’) (61) and 806R (5’-GGACTACNVGGGTWTCTAAT-3’) (62). PCRs, barcoding, library preparation and Illumina MiSeq sequencing were performed by the Joint Microbiome Facility (Vienna, Austria). First-step PCRs were performed in triplicate (20 μl vol per reaction) with the following conditions: 1X DreamTaq Buffer (Thermo Fisher), 2 mM MgCl2 (Thermo Fisher), 0.2 mM dNTP mix (Thermo Fisher), 0.2 µM of forward and reverse primer each, 0.08 mg.ml^-1^ Bovine Serum Albumin (Thermo Fisher), 0.02 U Dream Taq Polymerase (Thermo Fisher), and 0.5 µl of DNA template. Conditions for thermal cycling were: 95°C for 3 min, followed by 30 cycles of 30 sec at 95°C, 30 sec at 52°C and 50 sec at 72°C, and finally 10 min at 72°C. Triplicates were combined for barcoding (8 cycles). Barcoded samples were purified and normalized over a SequalPrep™ Normalization Plate Kit (Invitrogen), pooled, and concentrated on columns (Analytik Jena). Indexed sequencing libraries were prepared with the Illumina TruSeq Nano Kit as described previously (60), and sequenced in paired-end mode (2× 300 bp; v3 chemistry) on an Illumina MiSeq following the manufacturer’s instructions. The workflow systematically included four negative controls (PCR blanks, *i.e.,* PCR-grade water as template) for each 90 samples sequenced.

### Analysis of 16S rRNA gene amplicon sequences

Amplicon pools were extracted from the raw sequencing data using the FASTQ workflow in BaseSpace (Illumina) with default parameters. Input data was filtered for PhiX contamination with BBDuk (BBTools, Bushnell B, sourceforge.net/projects/bbmap). Demultiplexing was performed with the python package demultiplex (Laros JFJ, github.com/jfjlaros/demultiplex) allowing one mismatch for barcodes and two mismatches for linkers and primers. DADA2 (63) was used for demultiplexing amplicon sequencing variants (ASVs) using a previously described standard protocol (64). FASTQ reads 1 and 2 were trimmed at 220 nt and 150 nt with allowed expected errors of 2. Taxonomy was assigned to 16S rRNA gene/transcript sequences based on SILVA taxonomy (release 138). Sequencing of a nucleic acid extraction control and PBS used to collect the sorted cells in the FACS yielded 4 and 17 reads, respectively. Amplicon sequence libraries were analyzed using the vegan (v2.4.3) and phyloseq (v1.30.0) packages of the software R (https://www.r-project.org/, R 3.4.0). DESeq2 (42)(v1.26.0) implemented in phyloseq was used to determine statistically significant differences in ASV abundances between ATTO488^+^ and ATTO488^-^ sorted gates. Only ASVs that had ≥10 reads were considered for comparisons by DESeq2 analyses.

### *B. thetaiotaomicron* supplementation with miRNAs

A single colony of *B. thetaiotaomicron* strain VPI-5482 (DSM2079) was pre-grown in liquid *Bacteroides* Minimal Media (BMM) (65) under anaerobic conditions. The pre-grown culture was used to inoculate fresh BMM (1:100 dilution). The final suspension was aliquoted into a 96 well plate, and hsa-miR-21-5p MISSION® microRNA mimic, hsa-miR-21-5p scramble microRNA mimic and a double-stranded small RNA oligonucleotide control (sequence: 5’- GGAACGCCAACCGAAGUCUA - 3’) (all from Merck) were added to achieve a final concentration of 250 nM on each well. A water control was set up by adding nuclease-free water. Triplicate growths were established for each condition. The optical density (OD600nm) was measured every 30 minutes for a total of 18 hours on a Multiskan FC Microplate Reader (Thermo Scientific) placed inside the chamber. For transcriptomic analyses, the pre-grown culture was then taken at the exponential phase to inoculate pre-reduced BMM in glass tubes (6.6 mL per tube) and incubated at 37 °C until mid-exponential phase (OD600nm ∼ 0.5). MicroRNAs or a small RNA oligonucleotide control were added as above. A water control was set up by adding nuclease-free water. Triplicate growths were established for each condition. The cultures were then incubated with these small RNA molecules for one additional hour at 37°C, after which all tubes were removed from the chamber for subsequent fixation by the addition of 1:10 of the volume of a 2.5 mL acidic phenol/47.5 mL ethanol mixture. Samples were vortexed gently to mix, followed by a 5-minute incubation on ice. All samples were then transferred into 15 mL falcon tubes and centrifuged (3100 *g*, 4 °C) for 10 minutes and pellets were stored at -80 °C.

### *B. thetaiotaomicron* RNA isolation and purification

*B. thetaiotaomicron* culture pellets were defrosted on ice, homogenized in 100 μL of TE-Lyso buffer and incubated at 37°C for 15 minutes. The Norgen Biotek Total RNA Purification Kit (Norgen Biotek, Canada) was used to isolate and purify total RNA from the sample pellets, following the manufacturer’s instructions. The final RNA product was eluted in 41 μL nuclease-free water. DNase treatment (TURBO™ DNAse 2U/μL, Thermo Fischer Scientific) carried out according to the manufacturer’s instructions. Samples were incubated at 37°C with gentle shaking (250 rpm) for 30 minutes, after which an additional 1 μL of DNase was added to each tube and the incubation step was repeated (30 min, 37°C, 250 rpm). Following DNase treatment, the Zymo RNA Clean & Concentrator Kit (Zymo Research) was used to clean and purify the final RNA. The procedure for the extraction of large RNAs (> 200 nt) was followed, to eliminate any remaining small RNAs added as a treatment while retaining mRNAs. RNA product was eluted in 41 μL of nuclease-free water and the DNase treatment and RNA clean- up steps were repeated twice. The final RNA product was eluted in 24 μL nuclease-free water and stored at -80°C.

### RNA sequencing

For quality control and assessment of RNA integrity, automated electrophoresis was performed using the Agilent 4150 TapeStation system and RNA ScreenTape kit (Catalog number: 5067-5576, Agilent). One of the water triplicates had a low RNA Integrity Number equivalent (RINe) - 6.5, which is indicative of moderate RNA degradation. Because of this, only 2 water replicates were included in downstream analysis. The absence of any residual DNA contamination was confirmed via qPCR using the V4 16S rRNA gene-targeted primers (66, 67). The riboZero Plus Kit (Illumina) was used to deplete ribosomal RNA (rRNA) from the total RNA extracts, following the manufacturer’s protocol. Single-index barcoded sequencing libraries were prepared from rRNA-depleted RNA using the NEBNext® Ultra™ II Directional RNA Library Prep Kit for Illumina (NewEngland Biolabs) following the manufacturer’s protocol. Sequencing libraries were then pooled and sequenced on a HiSeq3000 (Illumina) in paired-end mode (150 cycles, 2x 75 bp reads) at the Biomedical Sequencing Facility (BSF) of the CeMM Research Center for Molecular Medicine of the Austrian Academy of Sciences/Joint Microbiome Facility (JMF) of the Medical University of Vienna and the University of Vienna (project ID JMF-2212-04).

Following sequencing, reads were quality-filtered and trimmed and mapped to the provided *B. thetaiotaomicron* reference genome (GCA_000011065.1, assembly ASM1106v1). Mapped read pairs were then counted using featureCounts (68, 69), and a subsequent differential expression analysis was performed using the DESeq2 package (v 1.26.0) in the software R (v 4.2.2) to define the differences between the controls and small RNA-treated conditions. Databases including NCBI (www.ncbi.nlm.nih.gov), QuickGO (www.ebi.ac.uk/QuickGO) and InterPro (www.ebi.ac.uk/interpro/) were used to manually annotate differentially expressed genes identified in the differential expression analysis. Additional packages in the software in R, including ComplexHeatmap (v 2.14.0) were used to visualize the results of the differential expression analysis.

### Quantification of miR-21 and miR-21^scr^ in *B. thetaiotaomicron* cultures

Bacterial cultures were spun down, supernatants were collected and frozen at -75 °C. RNA isolation was performed as stated above and eluted RNA was eluted in 41 µL nuclease-free water. DNase treatment and RNA purification were performed as stated above for transcriptomics analysis, except that an RNA purification protocol for total RNA clean-up was used. Next, cDNA synthesis and qPCR for miR-21 or miR21^scr^ were performed as mentioned above and miR21 concentrations were calculated with the use of a standard curve.

### Targeted quantification of *B. thetaiotaomicron* transcripts by RT-qPCR

Purified RNA from *B. thetaiotaomicron* cultures supplemented with small RNAs (isolated as described above) was reverse transcribed using the High-Capacity cDNA Reverse Transcription Kit (Thermo Fisher Scientific), according to manufacturer’s instructions, using 200 ng of RNA input. For qPCR iQ™ SYBR Green Supermix (Bio- Rad) was used, according to the manufacturer’s instructions, with 1 µl 1:20 diluted cDNA as input and 0.4 µM of primer (**Table S2**). qPCR was run in triplicates on a real-time PCR detection system CFX96 (BioRad) with the following thermal cycling conditions: 95°C for 5 min, followed by 40 cycles of 15 sec at 95°C, 20 sec at 56°C and 30 sec at 72 °C. ΔΔCt-method (70) was used to calculate relative expression levels of the *trp*AFCDGEB operon using guanylate kinase (*gmk*) as a reference gene (**Table S7**).

## Data availability

The 16S rRNA gene sequences were deposited in the NCBI (www.ncbi.nlm.nih.gov) Sequence Read Archive (SRA) under project number PRJNA1137825. RNA-Seq data is also available at NCBI under the same project (PRJNA1137825).

## Acknowledgements

This work was supported by the Austrian Science Fund via a Young Independent Research Group Grant (ZK-57) to F.C.P. and C.V. We would like to thank Marc Mussman for support with fluorescent-activated cell sorting, and Jasmin Schwarz, and Gudrun Kohl from the Joint Microbiome Facility of the Medical University of Vienna and the University of Vienna for assisting with amplicon and RNA sequencing. The authors declare that they have no conflict of interest.

