## Supplementary Figures 1-3 for "Human-derived microRNA 21 regulates indole and L-tryptophan biosynthesis transcripts in a prominent gut symbiont"

**This file contains:** Supplementary Figures 1 to 3.

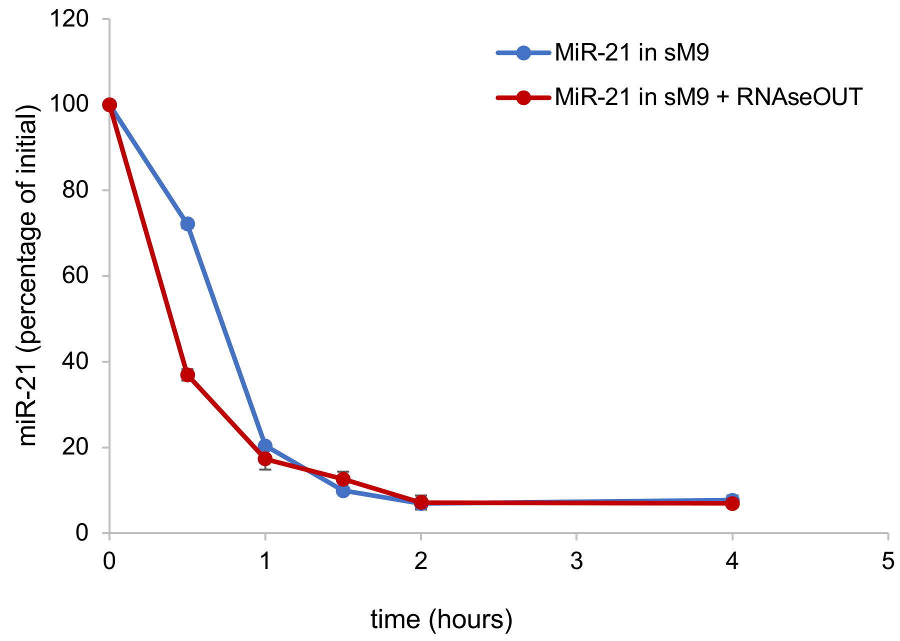


**Figure S1. Impact of RNAse treatments on miRNA association dynamics**. Percentage of the initial miRNA measured concentration remaining over time in live cell incubations with miR-21 alone versus miR-21 in combination with an RNase inhibitor (RNAseOUT). Data points represent the mean ± standard deviation of two replicates.

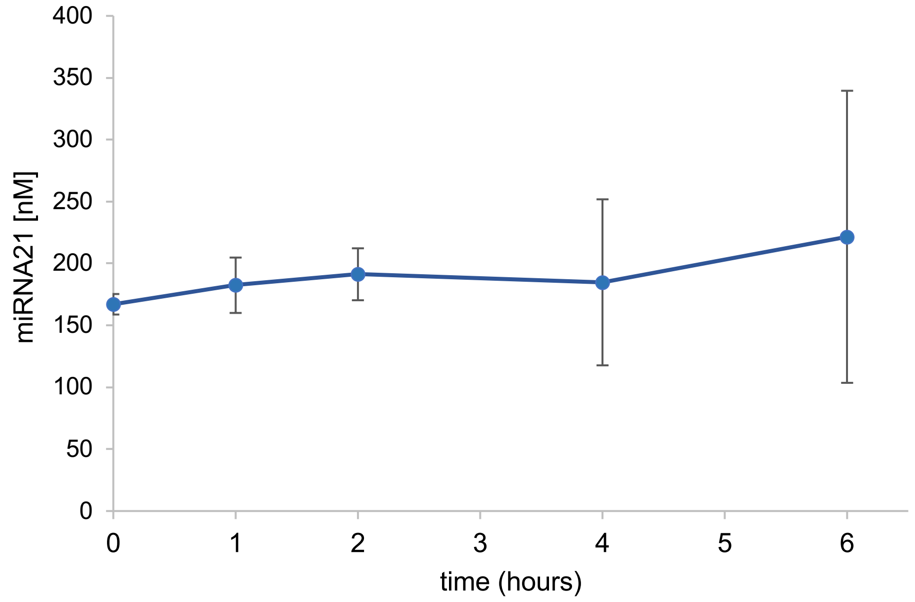


**Figure S2. Stability of miR-21 mimics in incubation vials**. Change in miR-21 concentration over time in incubation medium alone (sM9), as determined by qPCR. Data points represent the mean ± standard deviation of two replicates.

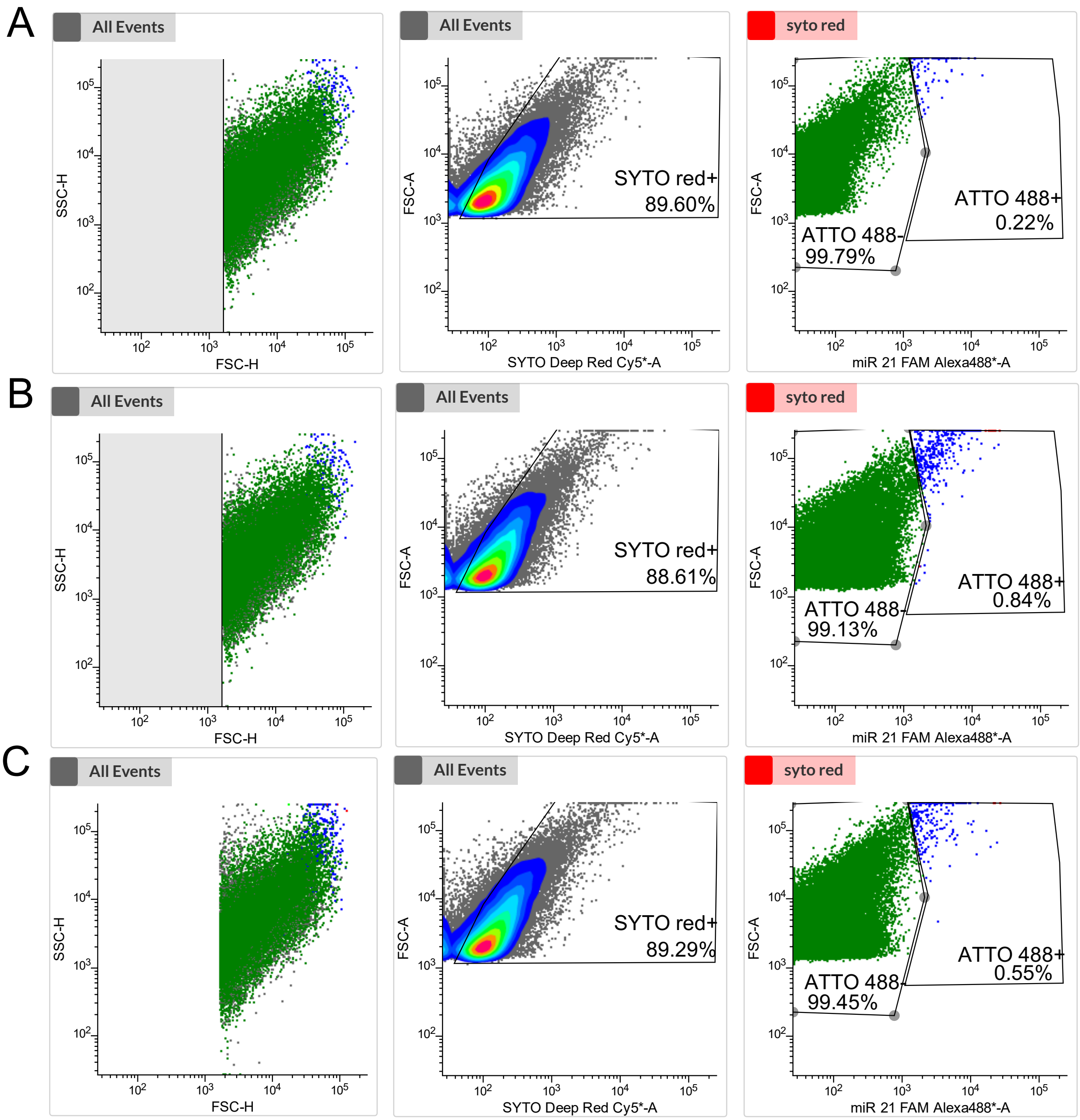


**Figure S3. Flow cytometry gating strategy to sort cell events interacting and not interacting with fluorescently labelled miRNAs.** A fixed staining-gating strategy to sort faecal sample cell events supplemented with water (**A**), ATTO488-labelled miR-21 (**B**), or ATTO488-labelled miR-21$^{scr}$ (**C**) is shown. Left panels show forward and side scatter plots with all events. Middle panels show a density plot with gating of SYTO red$^{+}$ cells (SYTO 62 red staining), to distinguish cells from debris and background noise. Percentages of SYTO red$^{+}$ events are indicated. Left panels show gating of ATTO 488$^{+}$ and ATTO 488$^{-}$ signal, which was performed using Alexa488 channel to discriminate cell events (SYTO red+) displaying or not ATTO 488$^{+}$ signal. The panels on the right also indicate the percentage of cells falling into the ATTO 488$^{+}$ and ATTO 488$^{-}$. The acquisition rate did not exceed 2,000 events per second. Each panel reflects over 100,000 registered events.
